# A method to estimate the frequency of chromosomal rearrangements induced by CRISPR/Cas9 multiplexing in Drosophila

**DOI:** 10.1101/815431

**Authors:** William A. Ng, Bruce H. Reed

## Abstract

Using CRISPR/Cas9 to simultaneously induce mutations in two or more target genes, commonly referred to as multiplexing, may result in chromosomal rearrangements such as inversions or translocations. While this may be undesirable in some contexts, the ability to recover chromosomal rearrangements targeted to specific sites in the genome is potentially a powerful tool. Before developing such tools, however, it is first important to measure the frequency with which chromosome rearrangements are induced by CRISPR/Cas9 multiplexing. To do this, we have developed a self-selecting screening system using a Drosophila line that carries an autosomal pericentric inversion in what is known as the autosynaptic form. All progeny of normal females crossed to males of this autosynaptic stock are lethal due to excessive aneuploidy. If an inversion is induced within the female germline, and if it is analogous to the inversion in the male autosynaptic line, then it is possible to recover progeny in which aneuploidy is reduced and viability is restored. Using this self-selection method, we screened 130 females and recovered one new autosynaptic element. Salivary gland polytene chromosome analysis, PCR, and sequencing confirmed the recovery of a breakpoint induced precisely between the two sgRNA target sites. Overall, we demonstrate that CRISPR/Cas9 multiplexing can induce chromosomal rearrangements in Drosophila. Also, in using this particular system, the recovery of chromosomal rearrangements was not a high frequency event.

## Introduction

The development of CRISPR/Cas9 genome editing has been a game-changing advance in the field of genetics. Co-expression of Cas9 with an engineered single guide RNA (sgRNA) forms a sequence homology-dependent endonuclease that creates DNA double-stranded breaks (DSBs) within a target sequence (1). DSBs induced by Cas9 are repaired by endogenous DNA repair machinery, either through error-prone non-homologous endjoining (NHEJ), or through homology directed repair (HDR). The CRISPR/Cas9 system is routinely used to facilitate gene knockouts and gene replacements in most model genetic organisms, including Drosophila (2–4).

At the heart of CRISPR/Cas9 is the localization of a protein/RNA complex to a specific sequence, as specified by a 17-20 nucleotide region of the sgRNA molecule. Examples of using CRISPR/Cas9 to engineer chromosome rearrangements in other model systems include the creation of an oncogenic translocation in mice (5), translocations in yeast (6), translocations and inversions in *C. elegans* (7, 8), and inversions in zebrafish (9). This study represents the first report of a chromosomal rearrangement induced by CRISPR/Cas9 in the Drosophila germline.

Site specific chromosomal rearrangements in Drosophila have been recovered using *FLP/FRT* mediated recombination (10). This technology was also used to generate a collection of chromosomal deletions with molecularly defined breakpoints, known as the DrosDel collection (11). Each of these cases, however, depended on random integration of a *P* element transposon carrying an *FRT* site. Subsequently, the design and recovery of deletions or rearrangements required knowing the genomic position and orientation of the insertions. The use of CRISPR/Cas9 multiplexing to generate chromosomal rearrangements would represent an improvement in that it can be used to target specific sequences and eliminates the need to map and characterize randomly generated insertions.

We wished to determine if CRISPR/Cas9 mutagenesis can be used to recover site specific chromosome rearrangements in Drosophila. To do this we developed a screening system based on autosynaptic chromosome elements (Fig.1). Due to the absence of meiotic recombination in Drosophila males, pericentric inversions, which are normally found in what is called their heterosynaptic form (*In(2LR)/+*, for example), can be maintained in what is known as the autosynaptic form (which is described as *LS(2)//DS(2)*) (12–14).

**Figure 1.**
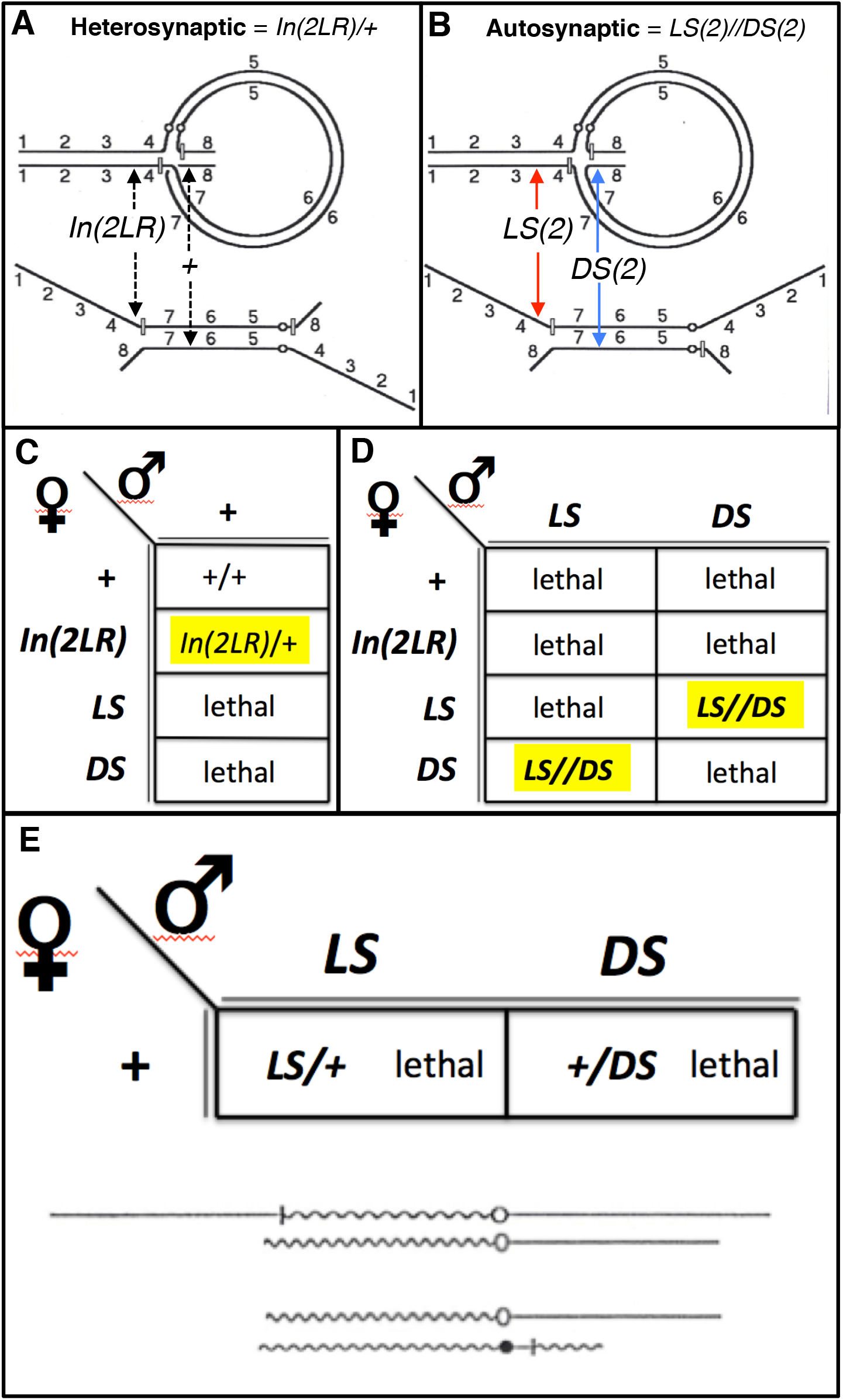
Autosynaptic Stocks: **(A)** Diagram showing the pairing arrangement of a normal chromosome paired with a pericentric inversion in the heterosynaptic form. Here the pericentric inversion occurs on a metacentric chromosome where one break is close to the centromere and the other is close to the telomere (upper diagram). Lower diagram shows an alternative way of depicting the same arrangement where pairing is only considered within the inversion loop. (**B**) Diagram showing the pairing arrangement of the same inversion but in the autosynaptic form (upper), and also showing an alternative way of depicting the same autosynaptic configuration (lower). From the lower diagrams it can be noted that autosynaptic stocks represent pericentric inversions where the two inversion breakpoints are no longer attached to a common centromere, but have been separated and are attached to the two homologous centromeres. (**C**) A Punnett square diagram showing the outcome of crossing a female that is heterozygous for a pericentric inversion crossed to a normal male. The heterozygous female can produce gametes that carry either the normal chromosome or the inversion, but can also produce gametes that carry the *LS* or *DS* autosynaptic chromosome elements as a result of a single exchange within the inversion loop. (**D**) A Punnett square diagram showing the outcome of crossing a female that is heterozygous for a pericentric inversion (or a female that carries the pericentric inversion in the autosynaptic form) crossed to a male carrying the inversion in the autosynaptic form. Females can produce gametes carrying the normal chromosome, the inversion, or either of the *LS* or *DS* autosynaptic chromosome elements. Due to the absence of crossing-over in male Drosophila, male gametes carry only the autosynaptic chromosome elements, and as a result the only viable progeny also carry the inversion in the autosynaptic form. (**E**) A Punnett square diagram showing the outcome of crossing a cytologically normal female to a male carrying the pericentic inversion it the autosynaptic form. In this example, progeny are *LS/+* (upper diagram), which carry an extra copy of the chromosome’s left arm and lack one copy of the distal region of the right arm, or *DS/+* (lower diagram) which lack one copy of the left arm and carry an extra copy of the distal region of the right arm. Both genotypes are lethal due to excessive aneuploidy.

We first recovered the autosynaptic form of a pericentric inversion with breakpoints at the base of *2L* and distal *2R*. Crossing schemes were then designed to recover females that have Cas9 germline expression and ubiquitous expression of two sgRNAs targeted to the approximate positions of the pre-existing *In(2LR)* breakpoints. These females were then crossed to males of the autosynaptic stock. In a self-selecting screen, where most progeny are lethal due to excessive aneuploidy, the only progeny to successfully complete development are expected to carry new autosynaptic elements derived from the female germline. From 130 females, we recovered one new autosynaptic element. Polytene chromosome analysis, PCR, and direct PCR sequencing of the PCR product of the anticipated *39E*;*60C* breakpoint were all consistent with the inversion breakpoint being the product of CRISPR/Cas9 mutagenesis. Although we lack a definitive frequency for this event, we conclude that it is possible to recover chromosomal rearrangements via CRISPR/Cas9 mutagenesis of two target sequences. To the best of out knowledge, this represents the first documentation of a chromosomal rearrangement recovered by targeted CRISPR/Cas9 mediated mutagenesis in Drosophila.

## Results

Many stocks carrying autosynaptic chromosome elements (Fig. 1) were recovered throughout the 1980’s; at the time, they were considered useful for synthesis of duplications and deficiencies (13, 14). The development of *FLP/FRT* based methods for recovering deletions and rearrangements was a vast improvement over methods based on autosynaptic chromosome elements. Being obsolete as a technology, all autosynaptic stocks were discarded from the main stock collection centers (Bloomington Drosophila Stock Center and Kyoto Stock Center). We were able to recover a stock of *In(2LR)lt*^*G16[L]*^*bw*^*v32g[R]*^ in the autosynaptic form using the product mimic method (12) (Fig. 2). *In(2LR)lt*^*G16[L]*^*bw*^*v32g[R]*^ has breakpoints in *2Lhet;60E* and *2Rhet;59E* and is a synthetic duplication of *59E-60E*; it was originally made using autosynaptic chromosome elements (15). Reciprocal crosses (females but not males can be outcrossed to lines with cytologically normal chromosomes) and polytene chromosome analysis were used to confirm that the stock was a bona fide *LS(2)//DS(2)* autosynaptic form of the *In(2LR)* (Fig. 3).

**Figure 2.**
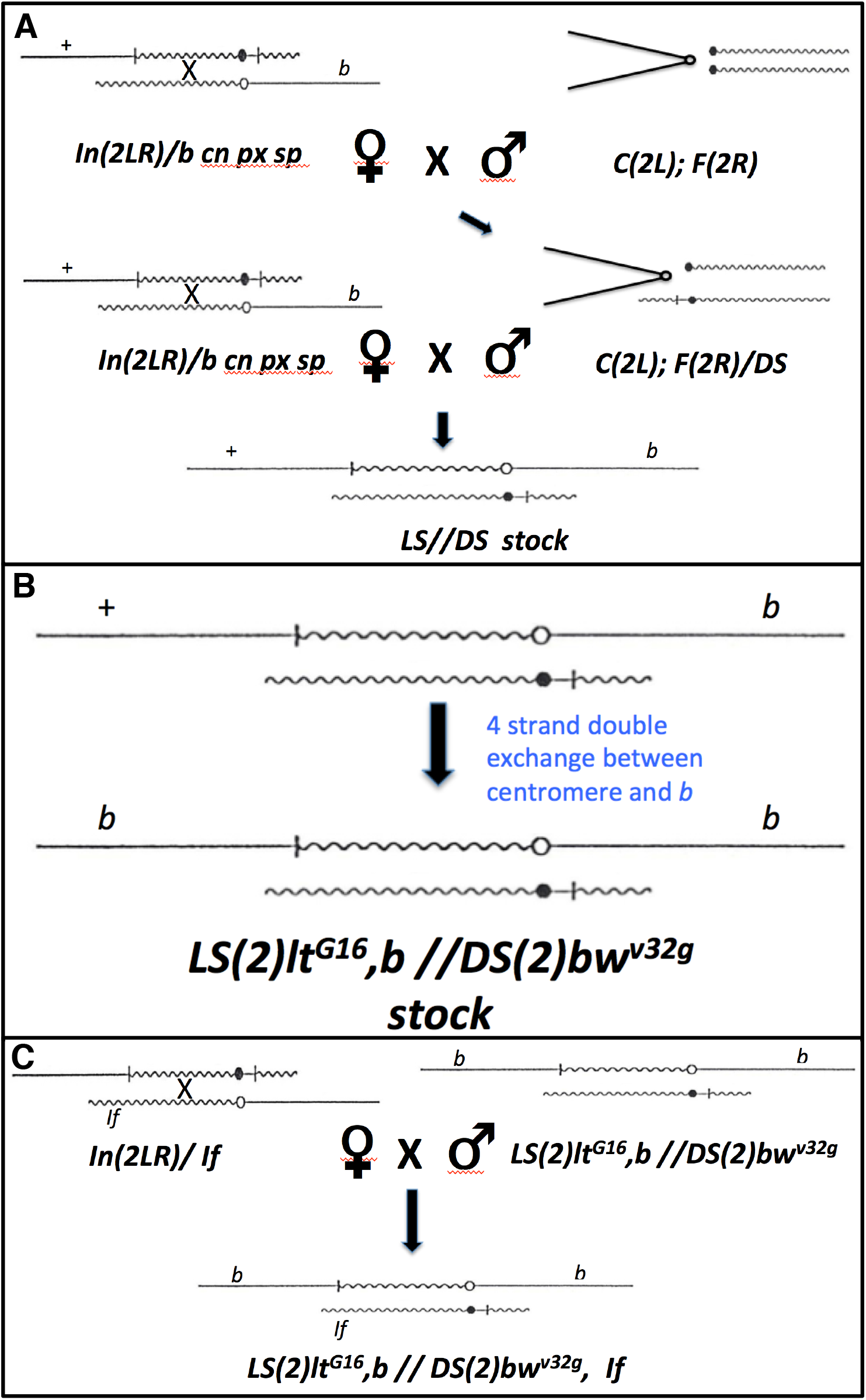
Recovery of the autosynaptic form of *In(2LR)lt*^*G16[L*^*]bw*^*v32g[R]*^ via the product mimic method (12): (**A**) The first step in recovering the autosynaptic form of *In(2LR)lt*^*G16[L]*^*bw*^*v32g[R]*^, which has breakpoints in *2Lhet;60E* and *2Lhet; 59E* and also is associated with a duplication of the region *59E-60E*, involves crossing females that are heterozygous for the inversion and a cytologically normal chromosome (here carrying the markers *b cn px sp*) to males of that carry a compound *2L* chromosome with free *2R* chromosomes. The same females are crossed with the male progeny of the first cross, where the males carry the compound *2L* chromosome with one free *2R* chromosome and the *DS(2)bw*^*v32g*^ autosynaptic chromosome element. The duplication of the distal *2R* region from the *59E* breakpoint to the telomere of *2R* represents a viable and tolerated degree of hyperploidy. Progeny of this cross are selected that carry the *LS* and *DS* elements. Males and females carrying *LS//DS* are crossed to each other to create the autosynaptic stock that will behave according to the Punnett square diagram as shown in Fig. 1D. (**B**) The *LS* element as recovered is heterozygous for the visible mutation *b*, but *LS* elements can become homozygous for *b* following a 4 strand double exchange between the centromere and *b* and can be selected within the *LS//DS* stock. (**C**) The original heterozygous inversion can be crossed to other visible markers, here shown is the dominant marker *Irregular facets* (*If*), and subsequently crossed to the *LS//DS* autosynaptic males to introduce markers onto the autosynaptic chromosome elements. Shown is the scheme used to recover *DS(2)bw*^*v32g*^ carrying the visible dominant marker *If*.

**Figure 3.**
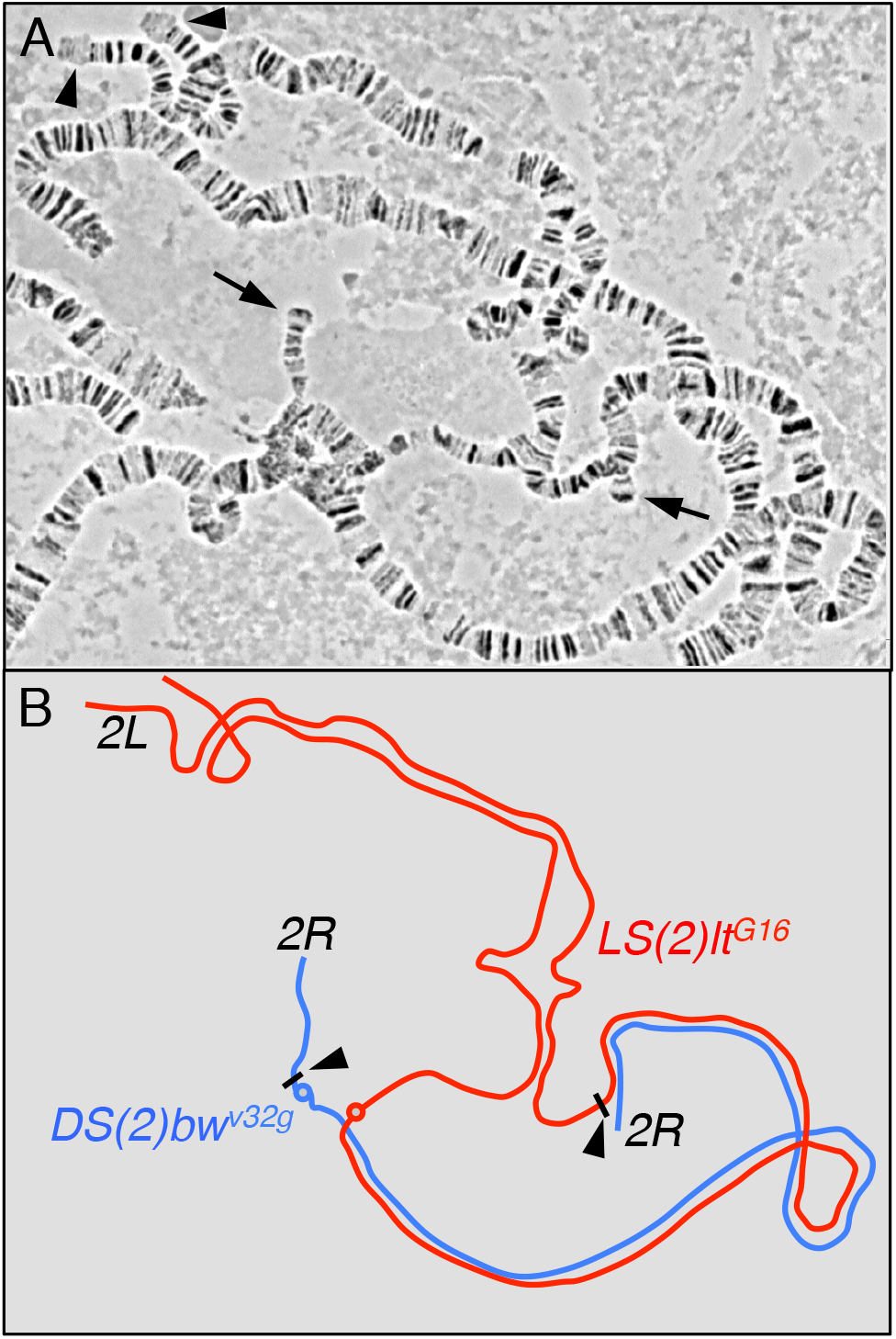
Cytological confirmation of the recovery of *LS(2)lt*^*G16*^ // *DS(2)bw*^*v32g*^: (**A**) Polytene chromosome squashes showing the inversion and chromosome breakpoints associated with *LS(2)lt*^*G16*^ // *DS(2)bw*^*v32g*^. In this example, the telomeres of *2R* (arrows) are asynapsed and the unpaired telomere of *DS(2)bw*^*v32g*^ is seen as separate and connected to the chromocenter. Most of *2L* (*2L* telomeres are indicated by arrowheads) and the entire inversion loop region are synapsed. (**B**) Schematic diagram showing the arrangement of the pairing of the *LS(2)lt*^*G16*^ (red) and *DS(2)bw*^*v32g*^ (blue) elements with the approximate positions of the breakpoints for each element indicated (black arrowheads).

Knowing the cytological breakpoints of *In(2LR)lt*^*G16[L]*^*bw*^*v32g[R]*^, we searched the Bloomington Drosophila Stock Center for one sgRNA line with a target slightly distal to *2Lhet* and a second sgRNA line having a target between *59E* and *60E*. These target sites were chosen such that any combination of *LS(2)lt*^*G16*^*//DS(2)*new\** or *LS(2)*new\**//*DS(2)bw*^*v32*^ would be slightly hyperploid would also complement any mutation induced at the breakpoint target site. We chose *TOE.GS00435*, which targets the region upstream of *CG2201* (map position *39E3*) and *TKO.GS00793*, which targets *cN-IIIB* (map position *60C2*).

A series of crosses culminated in the recovery of virgin females carrying *nos-Cas9*, which expresses *Cas9* in the germline, together with *TOE.GS00435* and *TKO.GS00793*, which ubiquitously express the sgRNAs targeted to 39E3 and 60C2, respectively (see Materials and Methods and Fig. 4 for description of crossing scheme). These females were mated in small groups (5-6 per vial) to autosynaptic males *LS(2)lt*^*G16*^//*DS(2)bw*^*v32g*^. As expected, most crosses were non-productive and produced no adult progeny. One vial, however, produced numerous progeny and virgin females isolated from this culture were found to be fertile upon backcrossing to the autosynaptic stock – which is a confirmation that they carry autosynaptic chromosome elements complementary to *LS(2)lt*^*G16*^ and *DS(2)bw*^*v32g*^. The segregation of visible markers allowed us to determine that these females carried a new *DS(2)* autosynaptic chromosome element, which we call *DS(2)CIA-1* (for <u>C</u>RISPR **<u>i</u>**nduced <u>a</u>utosynaptic). None of the progeny, however, were consistent with recovery of a new *LS(2)* element. Polytene squash analysis of *LS(2)lt*^*G16*^//*DS(2)CIA-1* clearly revealed a breakpoint between 39E3 and 60C2 (Fig. 5). PCR primers designed to amplify regions surrounding the sgRNA target sequences of *TOE.GS00435* and *TKO.GS00793* were tested in different combinations using template DNA from control or *LS(2)lt*^*G16*^//*DS(2)CIA-1*. As expected, forward and reverse primers for each target sequence generated product. Strikingly, we found a PCR product was generated using the forward primer for the *TOE.GS00435* target sequence with the reverse primer for the *TKO.GS00793* target sequence on template DNA from *LS(2)lt*^*G16*^//*DS(2)CIA-1*. Control DNA (from *LS(2)lt*^*G16*^//*DS(2)bw*^*v32g*^) did not generate a PCR product, and these results are consistent with a chromosomal breakpoint from one sgRNA target sequence to the other. This was confirmed by direct PCR sequencing (Fig. 6).

**Figure 4.**
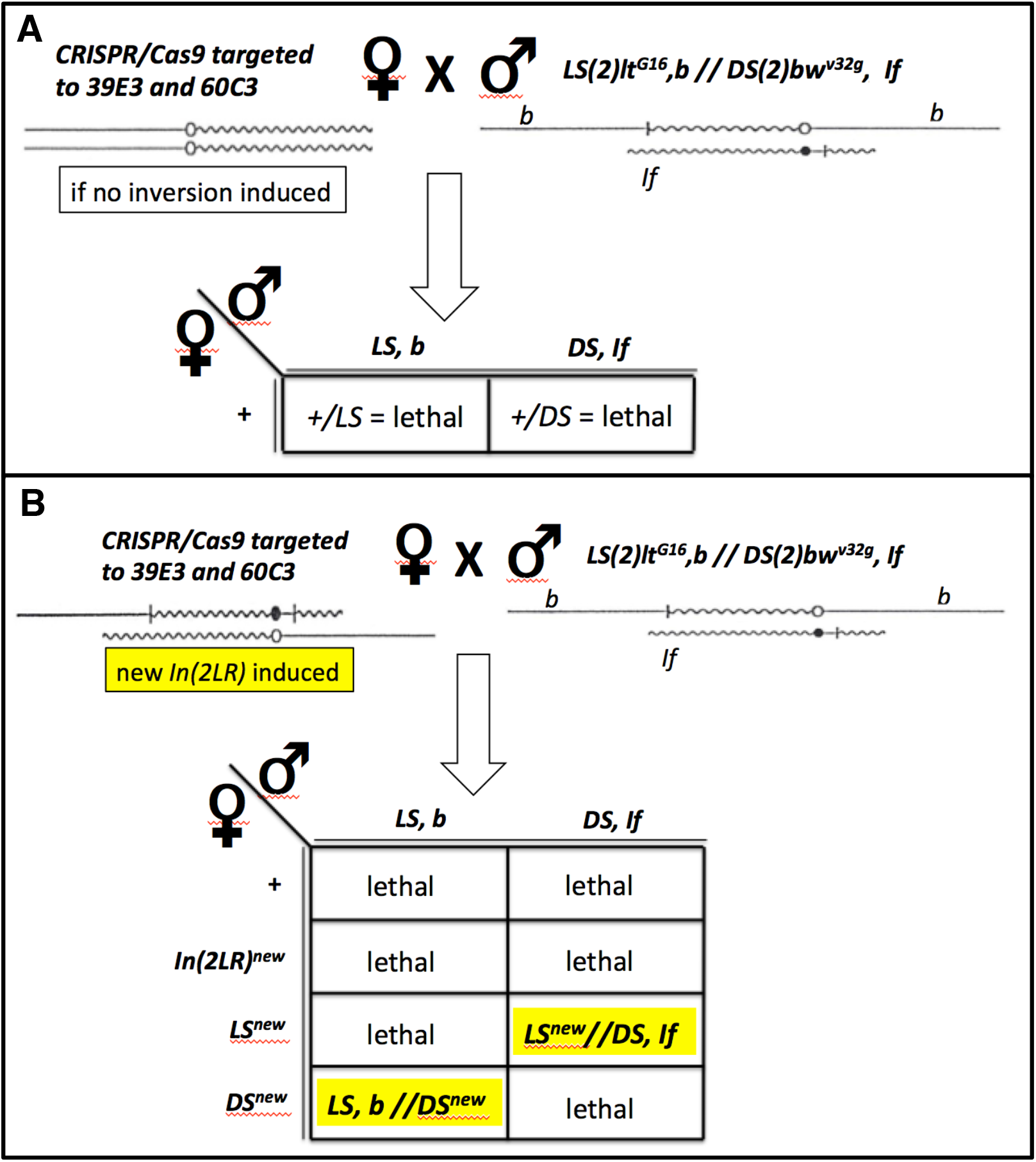
Autosynaptic chromosome elements can be used to design self-selecting screens: (**A**) Outcome of a cross of females carrying cytologically normal chromosomes that were exposed to active Cas9 and sgRNAs targeted to *39E3* and *60C2*. In the event that no inversion was induced, all progeny are lethal due to excessive aneuploidy. (**B**) Outcome of the same cross in which a new inversion was induced between the two sgRNA target sequences. The only viable progeny will carry new *LS* and *DS* elements derived from single exchanges within the inversion loop of a female harbouring a pericentric inversion within her germline.

**Figure 5.**
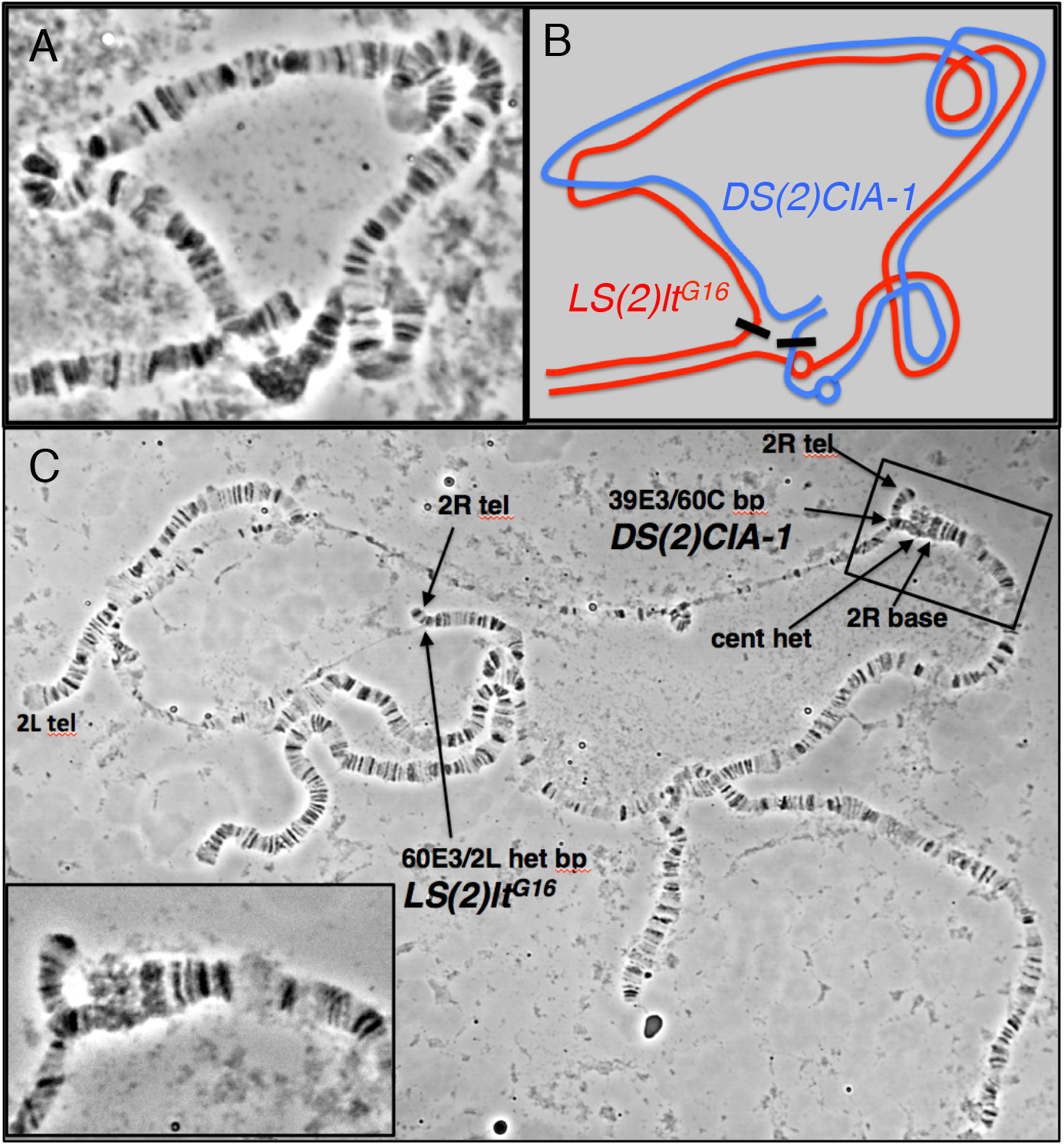
Cytological confirmation of the recovery of *LS(2)lt^G16^ // DS(2)CIA-1*: (**A**) Polytene chromosome squashes showing the inversion and chromosome breakpoints associated with *LS(2)lt*^*G16*^ // *DS(2)CIA-1*. In this example the homologous regions of the two elements have remained paired. (B) Schematic diagram showing the arrangement of the pairing of the *LS(2)lt*^*G16*^ (red) and *DS(2)CIA-1* (blue) elements with the approximate positions of the breakpoints for each element indicated (black arrowheads). (C) Example of a polytene squash of *LS(2)lt^G16^ // DS(2)CIA-1*where the *2R* telomeres are asynapsed. The base of *LS(2)lt*^*G16*^ and the tip of *DS(2)CIA-1* are indicated. The enlarged inset shows the *2R* telomere and the *39E;60C2* breakpoint associated with *DS(2)CIA-1*.

**Figure 6.**
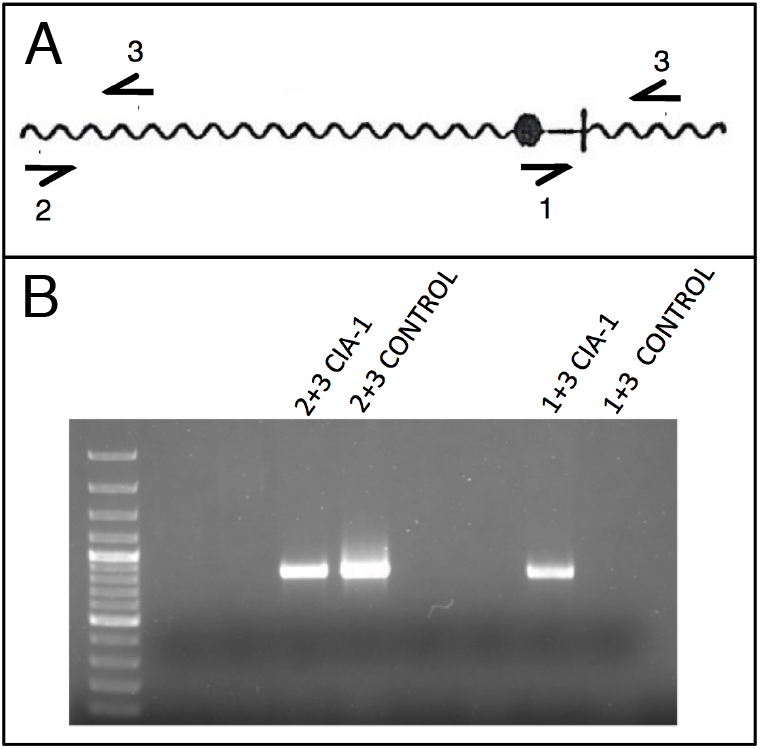
Molecular characterization of *DS(2)CIA-1*: (**A**) Overview of PCR strategy showing relative positions of primers designed to amplify the *60C2* sgRNA target sequence of *GS00793* (primers 2 + 3) and the *39E3;60C2* breakpoint using a primer flanking the proximal target sequence of sgRNA *GS00435* (primers 1 + 3). (**B**) Results of PCR showing products for primers 2+3 for in control (*LS(2)lt*^*G16*^*//DS(2)bw*^*v32g*^) and the new autosynaptic stock (*LS(2)lt*^*G16*^*//DS(2)CIA-1*) and a product for primers 1+3 only for the new autosynaptic stock and not the control.

**Figure 7.**
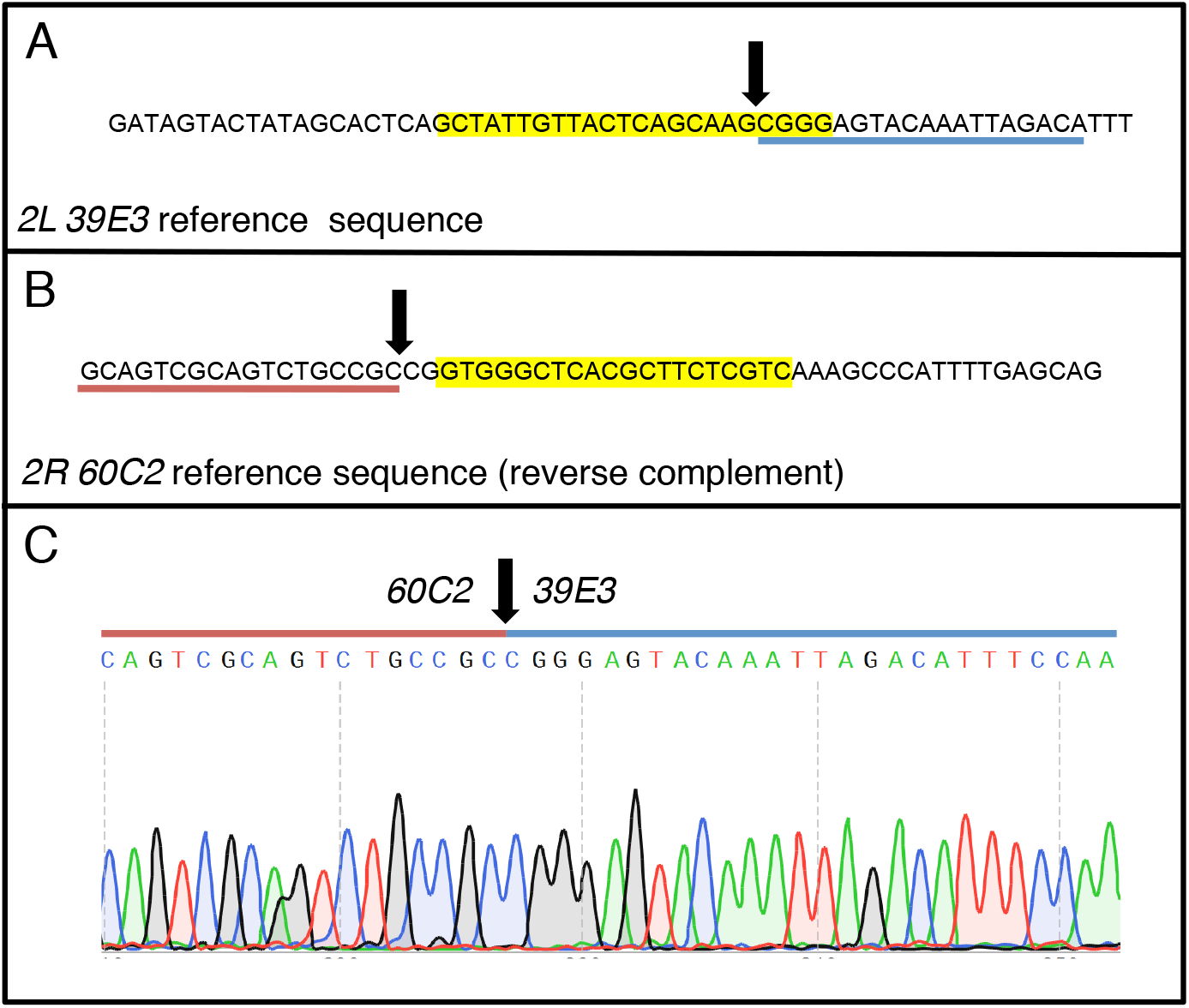
Sequence of *DS(2)CIA-1* breakpoint: (**A**) Reference sequence from region *39E3* showing the proximal sgRNA target sequence for *TOE.GS00435* (yellow highlight), the breakpoint position (arrow), and the region of sequence amplified using PCR primers 1+3 as show in Fig. 6 (blue underline). (**B**) Reverse complement of the reference sequence from region *60C2* showing the sgRNA target sequence for TKO.GS00793 (yellow highlight), the breakpoint position (arrow), and the region of sequence amplified using primers 1+3 as show in Fig. 6 (red underline). (**C**) Gas chromatogram DNA sequence readout generated by direct PCR sequencing of the primer 1+3 product, sequenced using primer 3. The molecular site of the *39E3;60C2* breakpoint of *DS(2)CIA-1* is indicated (arrow), and sequence from *60C2* (overscored by red line) is shown as contiguous with sequence from *39E3* (overscored by blue line).

## Discussion

We have reestablished autosynaptic stocks in order to develop a method for determining if CRISPR/Cas9 multiplexing can lead to chromosome rearrangements, and to measure the frequency of such events. In what was essentially a small-scale pilot screen, we targeted two regions of chromosome *2* by Cas9 mutagenesis within the female germline. The germline of such females could potentially harbor a pericentric inversion *In(2LR)* with breakpoints at *39E3* and *60C2*. The screening method to recover the pericentric inversion is self-selecting. This is because a cross of females with cytologically normal chromosomes mated to males carrying an analogous pericentric inversion in the autosynaptic form is non-productive; all progeny of such a cross are non-viable due to excessive eneuploidy. If, however, the germline of a female harbours a pericentric inversion as described, this female will segregate complementary autosynaptic chromosome elements following a single recombination event within the inversion loop. Our screening strategy was successful and we were able to recover a chromosome rearrangement induced by CRISPR/Cas9 mutagenesis.

In theory, it should be possible to recover both the *LS* and *DS* elements from the newly induced pericentric inversion. In our case, although we were able to demonstrate, for the first time in Drosophila, a targeted chromosome rearrangement using CRISPR/Cas9, we only recovered the *DS(2)* element. Similar observations have been made in self-selecting autosynaptic screens that were performed using X-ray mutagenesis (13). The explanation is found in the preferential and non-Mendelian inheritance of the smaller element during the segregation of asymmetric dyads in female Drosophila. In our case, a single recombination event within the pericentric inversion generates dyads that greatly differ in size (roughly an entire chromosome arm). Here the smaller *DS(2)* element is likely preferentially inherited by the oocyte nucleus while the larger *LS(2)* element is preferential passed to a polar body.

The screen as presented was performed on a small scale as we only tested 130 females. By repeating this screen we will be able to determine the frequency with which chromosomal rearrangements are recovered, at least in the context of our particular screen design. It is clear, however, that the recovery of progeny, and hence the induction of pericentric inversions within the female germline, was not a high frequency event.

The possibility of using CRISPR/Cas9 to design and generate chromosome rearrangements could be useful for studies regarding pairing dependent transcriptional regulation. Here, translocations or inversions could be targeted to precise sequences in order to test and map pairing dependent regulatory regions for a gene-of-interest. If the efficiency of CRISPR/Cas9 proves sufficiently high, or could be enhanced, then it may be also be possible to design schemes to recover chromosome rearrangements in other Drosophila, and perhaps non-Drosophila species. Overall, our self-selecting screen represents a platform in which the recovery of chromosome rearrangements induced by CRISPR/Cas9 can be rigorously tested and optimized.

## Materials and Methods

### Drosophila stocks

All Drosophila cultures were raised on standard medium at 25°C under a 12 hour light/dark cycle regime, unless otherwise indicated. The following stocks were obtained from The Bloomington Drosophila Stock Center:

#36569: *Df(2R)or*^*BR-6*^, *cn bw sp* / *In(2LR)lt*^*G16[L]*^*bw*^*v32g[R]*^

#68051: *y sc v sev^21^; P{y[+t7.7]v[+t1.8]=TOE.GS00435}attP40*

#77019: *y sc v sev^21^; P{y[+t7.7]v[+t1.8]=TKO.GS00793}attP40*

#54591: *y M{w[+mC]=nos-Cas9.P}ZH-2A w\**

The following stocks were obtained from The Kyoto Stock Center:

#101863: *y; C(2L)RM-SH1 / F(2R)VH2, bw*

#116105: *C(2L)RM-P2, dp^ov1^ / F(2R)1, cn c bw*

The following stock is maintained in the Reed Lab Stock Collection:

*b cn px sp*

### Recovery of the autosynaptic form of *In(2LR)lt*^*G16[L]*^*bw*^*v32g[R]*^

(Also see Fig. 2.) Females of *In(2LR)lt*^*G16[L]*^*bw*^*v32g[R]*^ / *b cn px sp* were recovered and crossed to males of *y; C(2L)RM-SH1 / F(2R)VH2, bw.* Male progeny that were *C(2L)RM-SH1 / F(2R)VH2, bw / DS(2)bw*^*v32g*^ were recovered and backcrossed to *In(2LR)lt*^*G16[L]*^*bw*^*v32g[R]*^/ *b cn px sp.* Progeny of this cross were established as individual lines and those that produced *b* progeny were selected as the autosynaptic stock *LS(2)lt*^*G16*^, *b* // *DS(2)bw*^*v32g*^. Females of *In(2LR)lt*^*G16[L]*^*bw*^*v32g[R]*^ / *If* were then crossed to *LS(2)lt*^*G16*^, *b* // *DS(2)bw*^*v32g*^ males, permitting recovery of *LS(2)lt*^*G16*^, *b* // *DS(2)bw*^*v32g*^, *If*.

### Screen for new autosynaptic elements induced by CRISPR/Cas9

Males were recovered that were *w nos-Cas9.P w / Y; TOE.GS00435 / +* and these were crossed to *FM7 / +; CyO / +* females. Virgin female progeny of this cross that were *FM7h,w / y nos-Cas9.P; CyO* / *TOE.GS00435* were collected and crossed to males of *y sc v sev*^*21*^*; TKO.GS00793.* Virgin females of this cross that were *y nos-Cas9.P / y sc v sev*^*21*^*; TOE.GS00435/ TKO.GS00793* were collected and crossed to *LS(2)lt*^*G16*^,, *b* // *DS(2)bw*^*v32g*^, *If* males. Rare progeny were backcrossed to *LS(2)lt*^*G16*^,, *b* // *DS(2)bw*^*v32g*^, *If* males. Viable progeny from this cross indicated the recovery of a new autosynaptic stock. Progeny that were *b If*^+^ indicated a new *DS(2)* element while *b^+^ If* progeny indicated recovery of a new *LS(2)* element (Also see Fig. 4).

### Polytene Chromosome Squashes

Temporary salivary gland polytene chromosome squashes were prepared using a 2% natural orcein stain (Gurr’s 23282) that was prepared in equal parts 45% acetic and 45% lactic acid and aged for 29 years. Polytene chromosome breakpoints were mapped with reference to Lefevre’s photographic maps (16).

### Molecular characterization of *DS(2)CIA-1*

In order to molecularly characterize the *DS(2)CIA-1* breakpoint we extracted genomic DNA from a pooled group of 20-30 isogenic male flies using gSYNC DNA Extraction Kits (Geneaid) according to supplier’s instructions. We used 2.5 μL of isolated genomic DNA in 25 μL PCR reactions (2X FroggaMix; FroggaBio) utilizing primers flanking the guide target regions at ~400 bp upstream and downstream of the sgRNA target sequences. PCR products were purified using GenepHlow Gel/PCR Kits (Geneaid) then sent out for Sanger sequencing using both forward and reverse primers.

## Acknowledgements

Stocks obtained from the Bloomington Drosophila Resource Center (NIH P40OD018537) were used in this study. This work was supported by a grant to B.H.R. from the Natural Sciences and Engineering Research Council of Canada (NSERC RGPIN-2015-04458).

